# Fucoidan exerts anti-cancer activity in a 3D prostate spheroid cell culture model

**DOI:** 10.1101/2025.11.18.689081

**Authors:** Jelena Vider, Todd Shelper, Katherine Ting Wei Lee, J. Helen Fitton, Nicholas Peter West

## Abstract

To investigate the effect of fucoidan on the viability, migration and mRNA/microRNA profiles of two prostate cancer cell lines in 3D culture (spheroids). Hormone resistant, PC-3, and hormone sensitive, LNCaP, spheroids were incubated in the presence of a proprietary fucoidan blend extract from *Undaria pinnatifida* and *Fucus vesiculosus*. Magnetic bioprinting was used to assess cells’ migration/metastatic potential in response to fucoidan exposure. Cellular viability was assessed using a cytotoxicity lactate dehydrogenase (LDH) assay. nCounter (Nanostring) assays were used for microRNA and mRNA profiling in control and treated cells. Fucoidan treated PC-3 spheroids clearly showed a dose dependent viability/migration response and changes in spheroid structural morphology. Fifty-seven genes were significantly differentially expressed between fucoidan treated and non-treated PC-3 spheroids. No miRNAs at an adjusted p-value were significantly differentially expressed in PC-3 spheroids with fucoidan co-culture. Thirty-two genes were significantly differentially expressed in the fucoidan treated to non-treated LNCaP spheroids and there was a significant 3.5-fold (pAdj=0.012) increase in hsa-miR-1246 in the fucoidan treated compared to non-treated LNCaP spheroids. Gene sets enriched in fucoidan treated PC-3 spheroids related to DNA repair processes, cellular development, cell cycle and inflammation. These findings provide further pre-clinical evidence for a role for fucoidan in targeting cancer related pathways in both hormone sensitive and hormone resistant prostate cancer.

## Introduction

Prostate cancer (PCa) is the second most prevalent cancer in men worldwide (1). Most cases of death from PCa are due to metastasis. Hormone sensitive prostate cancer is treatable with a variety of drugs and surgical procedures, with combinatorial therapy comprising hormone treatment or chemotherapy with androgen deprivation improving survival (2). However, hormone resistant prostate cancer has limited options for control and the prognosis for advanced cancer remains poor. The need for new therapeutic strategies is required to improve survival in individuals with prostate cancer.

Fucoidans are brown seaweed derived sulfated fucose rich polysaccharides that have been reported to modulate numerous biological functions. These include anti-inflammatory effects via suppression of pro-inflammatory cytokines (3, 4) and inhibition of adhesion of pathogens (5, 6). Fucoidan is also known as a selectin and scavenger receptor blocking agent (7), and to exert anti-cancer activity across a range of cancer types and models (4). Fucoidan derived from *Undaria pinnatifida* has been previously shown to cause selective cell cycle arrest *in vitro* in colon cancer cells (8), whilst primary fibroblasts were unaffected. Fucoidan has been demonstrated to inhibit prostate cancer cell line proliferation in monolayer culture i*n vitro* and in mice studies (9-12). Importantly, fucoidans have a demonstrated record of safety across various population groups (13-15).

Fucoidan extracts from *Undaria pinnatifida* and *Fucus vesiculosus* are commercially available for inclusion in dietary supplements, and natural components of the edible seaweed ‘wakame’ and ‘bladder wrack’, respectively. Dietary fucoidan has been shown to modulate innate immunity in various models (16), which may also be a route via which it exerts an anti-cancer effect *in vivo*. Although not examined here, indirect effects on gut, urine or tumour microbiome may be connected to both inflammation, immunity, and the progression of cancer (17, 18). In a clinical study, ingestion of fucoidan was shown to restore a marker of gut innate immunity (lysozyme) to normal levels in athletes (19), although the effects on gut microbiome are not described. Given the evidence of anti-cancer effects with dietary fucoidan *in vitro*, fucoidans may be a promising adjunct strategy in PCa.

LNCaP and PC-3 cells are two cell lines that respectively represent both hormone sensitive and hormone resistant prostate cancer (20). 3D culture is increasingly popular as a screening tool as it has superior correlation with *in-vivo* data as compared to standard 2D culture systems (21). Self-assembling 3D spheroid cultures can model tumoroid-like structures *in-vitro*, incorporating important physical properties, such as increased cell-cell interactions, and compound exposure gradients that are otherwise absent from monolayer cultures. Indeed, prostate cancer organoid cell culture has been reported to recapitulate the molecular diversity of prostate cancer and been successfully used in drug screening (22). The effect of fucoidan on 3D cultured prostate cancer cells has not previously been examined.

The aim of this research was to characterize the effect of a proprietary fucoidan blend that includes extracts from *Undaria pinnatifida* and *Fucus vesiculosus* on miRNA and mRNA expression in hormone sensitive and hormone resistant PCa cells. Transcriptional profiling in prostate cancer is improving PCa diagnosis and prognosis, and evidence that fucoidan exerts effects on dysregulated miRNA and mRNA expression within tumour cells may provide important new data for a naturally occurring seaweed compound to underpin translation to clinical research.

## Materials and Methods

### Cell culture

PC-3 (#90112714) and LNCaP (Clone FGC; # 89110211) cell lines from The European Collection of Authenticated Cell Cultures (ECACC) were obtained from Merck/ Sigma-Aldrich Pty Ltd.

PC-3 cell line was cultured in F12K media (ATCC^®^ 30-2004); LNCaP cell line was cultured in RPMI-1640 Medium (ATCC 30-2001). Cell lines were cultured with 2% fetal bovine serum (FBS) content in standard cell culture conditions (humidified incubator at 37°C, 5% CO_2_).

Elplasia 96-well plates (Corning #4442) were used to generate a high density of spheroids in a scaffold-free model according manufacturer’s protocol. Briefly, cells were seeded at 5x10^4^ cells per well. The formation of spheroids was recorded using EnSight imaging plate reader (PerkinElmer) and media was collected for the LDH assay.

Greiner Bio-One’s Magnetic 3D Cell Culture system was used for the reproducible formation of one spheroid per well in 96-well plate with Cell-Repellent surface, following manufacturer’s protocol. Briefly, cells were magnetized by overnight incubation with biocompatible NanoShuttle™-PL, trypsinized and then 5x10^4^ cells were forced by magnetic bioprinting to form structurally and biologically representative 3D tumoroid model *in-vitro*.

Fucoidan extract from *Undaria pinnatifida* and *Fucus vesiculosus* was blended and provided by Marinova Pty Ltd. The proprietary aqueous extract was designed for ingestion, with a standardised fucoidan content greater than 85% as determined by previously reported methods (3). A hydrolysed sample was assessed by the phenol-sulfuric technique from Dubois for total carbohydrate (23). The carbohydrate profile was determined using a GC-based method for the accurate determination of individual monosaccharide ratios in a sample. This method relies on the preparation of acetylated alditol derivatives of the hydrolyzed samples. The uronic acid content was determined by spectrophotometric analysis of the hydrolyzed compound in the presence of 3-phenylphenol, based on a method described by Filisetti-Cozzi and Carpita (24). Sulfate content was analyzed spectrophotometrically using a BaSO_4_ precipitation method (BaCl_2_ in gelatin), based on the work of Dodgson (25). Cations, including Na^+^, K^+^, Ca^2+^, and Mg^2+^, were determined by Flame Atomic Absorption Spectroscopy.

Fucoidan blend solution was prepared at final concentration of 400 µg/ml in cell culture media with 2% FBS. Cells were exposed to fucoidan blend concentrations up to 200 µg/mL for up to 144 hrs without media change.

### Cell viability

Count/viability MUSE assay (Merck) was used to assess cells count/viability prior to starting the experiments. EnSight imaging plate reader was used to assess the effect of fucoidan treatment on PC-3 and LNCaP cells in 3D culture conditions (data is not shown).

### Cytotoxicity assay

To assess fucoidan’s EC_50_ CytoTox 96^®^ Non-Radioactive Cytotoxicity Assay (Promega, #G1781) was used according to manufacturer’s protocol. Briefly, cells were seeded to Elplasia 96-well plates (Corning #4442) at 5x10^4^ cells per well and treated with serial dilutions of fucoidan blend for up to 144 hrs. Media for LDH assay was collected at 24, 48, 72, 96 and 144 hrs

### Live Imaging

The magnetic 3D cell culture system was visualized with a Sartorius Incucyte S3 Live-cell analysis system using a 4x objective. A single brightfield central FOV was acquired from every well of a Greiner ULA 96 well plate every 3 hours for a total of 138 hours.

### Spheroid Analysis

Analysis of spheroid growth and compound effect kinetics were assessed using a traditional user-defined image analysis pipeline developed in CellProfiler (Ver 4.2.1)(26). Briefly, raw brightfield TIFF images from an Incucyte were first flatfield corrected using the illumination correction module then the images were inverted. Next, using the ImageMath module images were first opened then closed with a final smoothing process to assist with brightfield segmentation algorithms. Finally, spheroids were segmented from background using the IdentifyPrimaryObjects module, filtered and multiple size/shape and texture features were measured. The 2D area measurement was used to assess changes in spheroid size and compactness.

### nanoString nCounter PanCancer Pathways

Total RNA was extracted using the Maxwell^®^ RSC miRNA from Tissue kit (Promega) and used for subsequent experiments on the NanoString nCounter platform (NanoString, Seattle, USA) according to the manufacturer’s guidelines. Gene expression was measured using the NanoString nCounter PanCancer Pathways Panel (NanoString, Seattle, USA) containing 730 genes and 40 housekeeping genes. Briefly, a total of 300ng of total RNA was hybridized for 20 hours and counted on the nCounter Digital Analyser (NanoString, Seattle, USA). Raw data was normalized by subtracting the mean plus one standard deviation of eight negative controls while technical variation was normalized through internal positive controls. Data was corrected for input volume via internal housekeeping genes. A threshold of 20 was set as the lower limit of detection.

### NanoString nCounter miRNA

miRNAs were measured using the commercially available nCounter Human v3 miRNA assay (NanoString, Seattle, USA) targeting 798 miRNAs and including positive and negative controls, housekeeping transcripts and ligation specific controls for quality control of sample integrity and platform error. Briefly, 300 ng RNA was hybridized for 20 hours and counted on the nCounter digital analyzer (NanoString, Seattle, USA). Normalization and background correction were undertaken using positive and negative spiked-in controls.

### nSolver/Rosalind software

mRNA and miRNA data analysis were performed using nSolver version 4.0 (NanoString, Seattle, USA) and Rosalind software. Following background correction and normalization, differential expression and pathway analysis between the conditions (fucoidan blend treated and non-treated) were completed in Rosalind ((https://rosalind.onramp.bio/).

### Statistics

Significance was set at *p*=0.05 using a Bonferroni correction for multiple comparisons. All data are expressed as mean ± standard deviation (SD) or mean ± 95% confidence interval (CI) unless otherwise stated.

## Results

Prostate cancer cell lines were cultured in a 3D cell culture system. Cell viability, migration, mRNA and miRNA analysis were carried out.

### Cell culture growth assays in 3D format

#### Cell viability

Cell viability in the presence of the blended fucoidan was measured by LDH release and differential effects were observed in relation to PC-3 and LNCaP cells. Increased LDH release was observed from PC3 cells at a concentration of 6.25 µg/ml, while LNCaP cell viability was only altered at a concentration of >100 µg/ml (Figure 1a and 1b). Data on LDH release by the cell lines individually is shown at Supplementary Figure (SF) 1.

**Figure 1.**
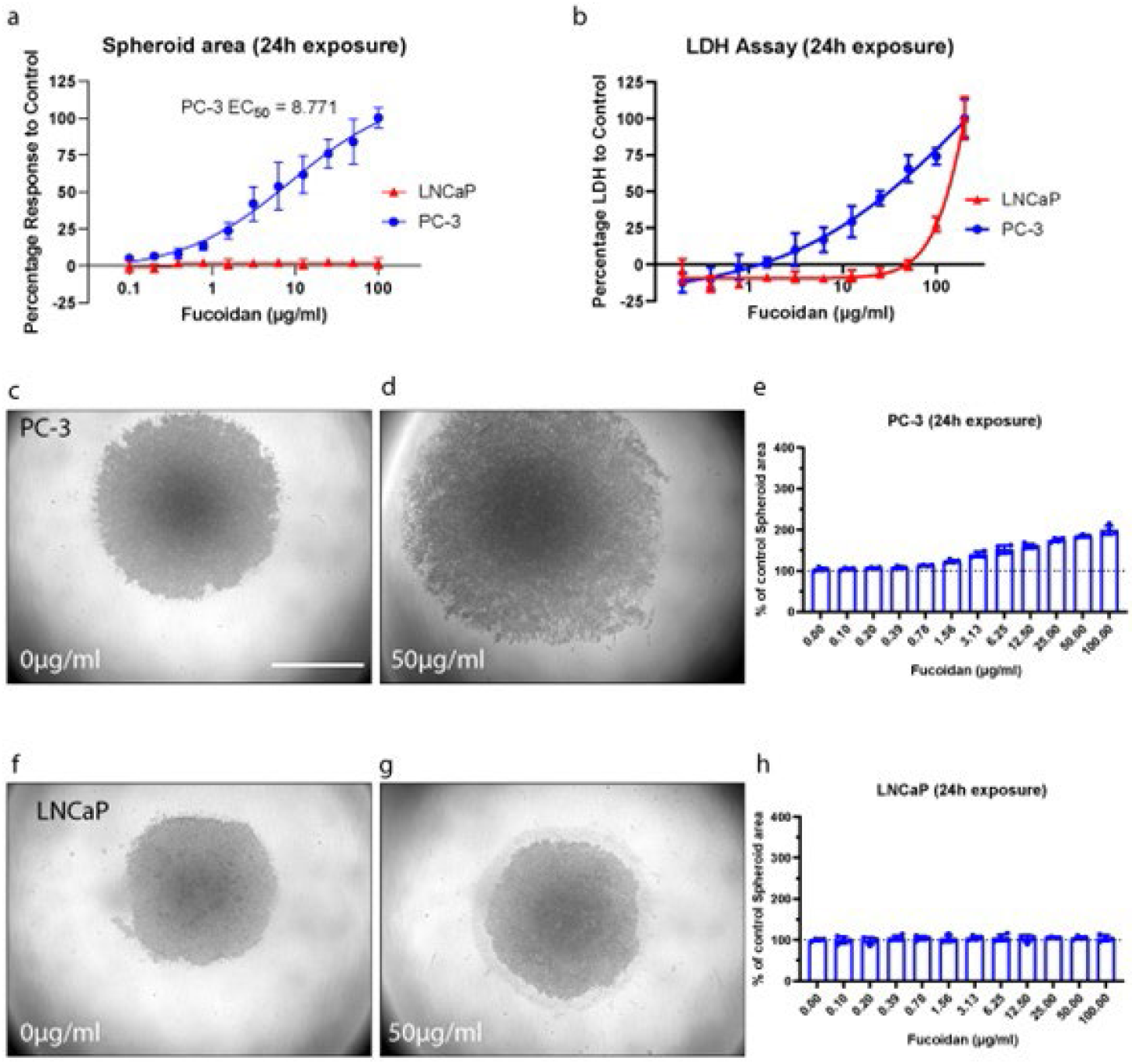
Spheroid area quantification and cellular cytotoxicity data from PC-3 and LNCaP 3D cultures. (a) Dose response curve of fucoidan blend plotted for LNCaP and PC-3 cell lines, area normalized to control well spheroid area at the 24h timepoint. (b) LDH release from PC-3 and LNCaP cells cultured in the presence of the fucoidan blend. Representative brightfield image of a PC-3 spheroid from the vehicle only control (c) and the 50µg/ml treatment group (d). Representative brightfield image of a LNCaP spheroid from the vehicle only control (f) and the 50µg/ml treatment group (g). Changes to the area occupied by PC-3 (e) and LNCaP (h) cells normalized to control well at the 24 timepoint. Scale bar = 1000µm. Brightfield images were flatfield corrected with linear contrast adjustments only. Data are represented as the average of quadruplicate spheroids with SD error bars.

#### 3D cell bioprinting

Following 15 minutes incubation in Greiner Bio-One’s Magnetic 3D Cell Culture system, both PC-3 and LNCaP cells self-aggregated to form tightly packed multicellular 3D structures by 24h (SF2). As seen in SF3, each cell type follows a distinct and reproducible growth profile. A dose dependent loss in cell-cell attachment, which resulted in a disintegration of spheroid structure was observed with fucoidan blend incubation for the PC-3 cell line above concentrations of 6.25 µg/ml. As can be seen in Figure 1d, cells dissociate from the solid spheroid structure to form loosely packed single cell layer on the bottom of the well. Fucoidan blend treatment displayed an EC_50_ of 8.77 µg/ml for this loss of cell attachment phenotype at 24h exposure (Figure 1a). At the 50 µg/ml dose an 84.3% (80.7 – 87.8) increase in total cell area was observed. (Figure 1e). LNCaP 3D cultures exposed to fucoidan did not display this phenotype, even at the top dose tested, with cells remaining tightly packed in the spheroid (Figure 1f-h). Additional EC_50_ time point data from PC-3 spheroid area measurement (24h – 96h) is shown in SF4. Additional timepoint spheroid area data for both PC-3 and LNCaP cell lines (24h – 96h) is shown in SF5.

### mRNA analysis

#### PC-3

Figure 2 displays differentially expressed genes between PC-3 spheroids treated with fucoidan and control. There were 38 genes that were significantly increased in expression and 19 genes that were significantly decreased in expression at an adjusted *p*-value<0.05 in the fucoidan treated PC-3 spheroids after 24h exposure (Supplementary Table 1). Unsupervised hierarchical clustering of the genes that were significantly differentially expressed revealed two clusters that corresponded to the treated and untreated cell lines (Figure 2A and B).

**Figure 2.**
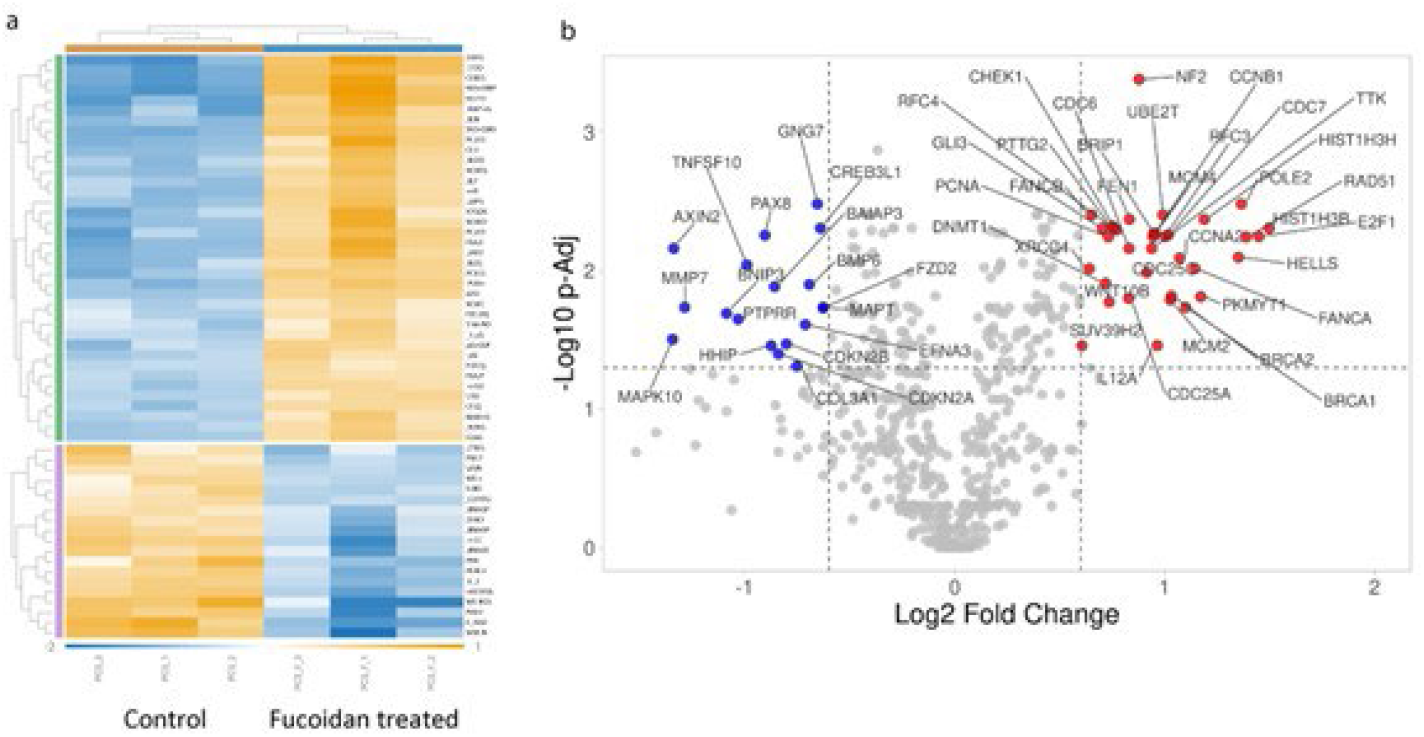
(a) A heatmap of differentially expressed genes between treated and untreated PC-3 spheroids. Each row is a gene and each column a sample. Yellow is increased expression and blue is decreased expression (b) Volcano plot showing differentially expressed genes between PC-3 spheroids treated with fucoidan (PC-3 F) v untreated cells (PC-3); blue-decreased, red-increased.

Gene Set Analysis (GSA) between the treated and untreated PC-3 cell lines indicates that treated cells were enriched in the cell cycle/apoptosis and the DNA Damage/repair pathways while the untreated cells were enriched in the PI3K-AKT, MAPK, RAS, Wnt and transcriptional misregulation (Figure 3).

**Figure 3.**
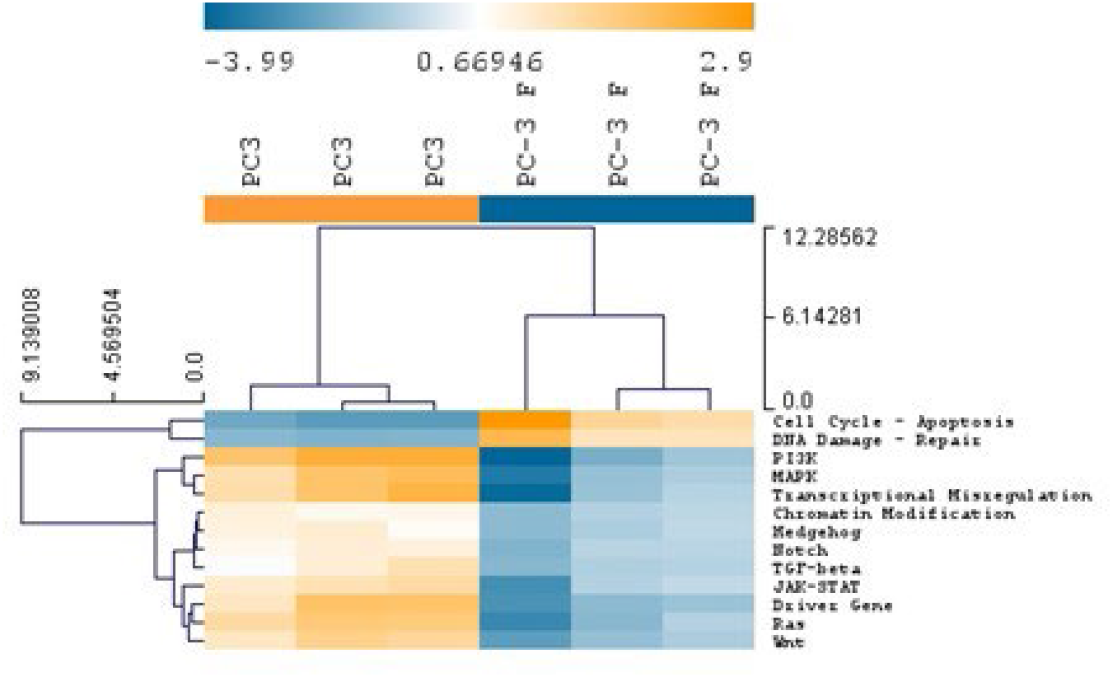
A heatmap of pathway scores for PC-3 spheroids treated with fucoidan and PC-3 spheroids not treated Fucoidan. Each column is a cell line and each row a pathway. Orange is increased pathway expression and blue is decreased pathway expression.

#### LNCaP

Figure 4 displays differentially expressed genes between LNCaP hormone dependent cancer spheroids treated with fucoidan and control. There were six genes that were significantly increased in expression, including HPGD a tumour suppressor, and 26 genes that were significanatly decreased in expression at an adjusted *p*-value in the fucoidan treated cells (Figure 4a and Supplementary Table 2). Unsupervised hierarchical clustering of the genes that were significantly differentially expressed revealed two clusters that corresponded to the treated and untreated spheroids (Figure 4b).

**Figure 4.**
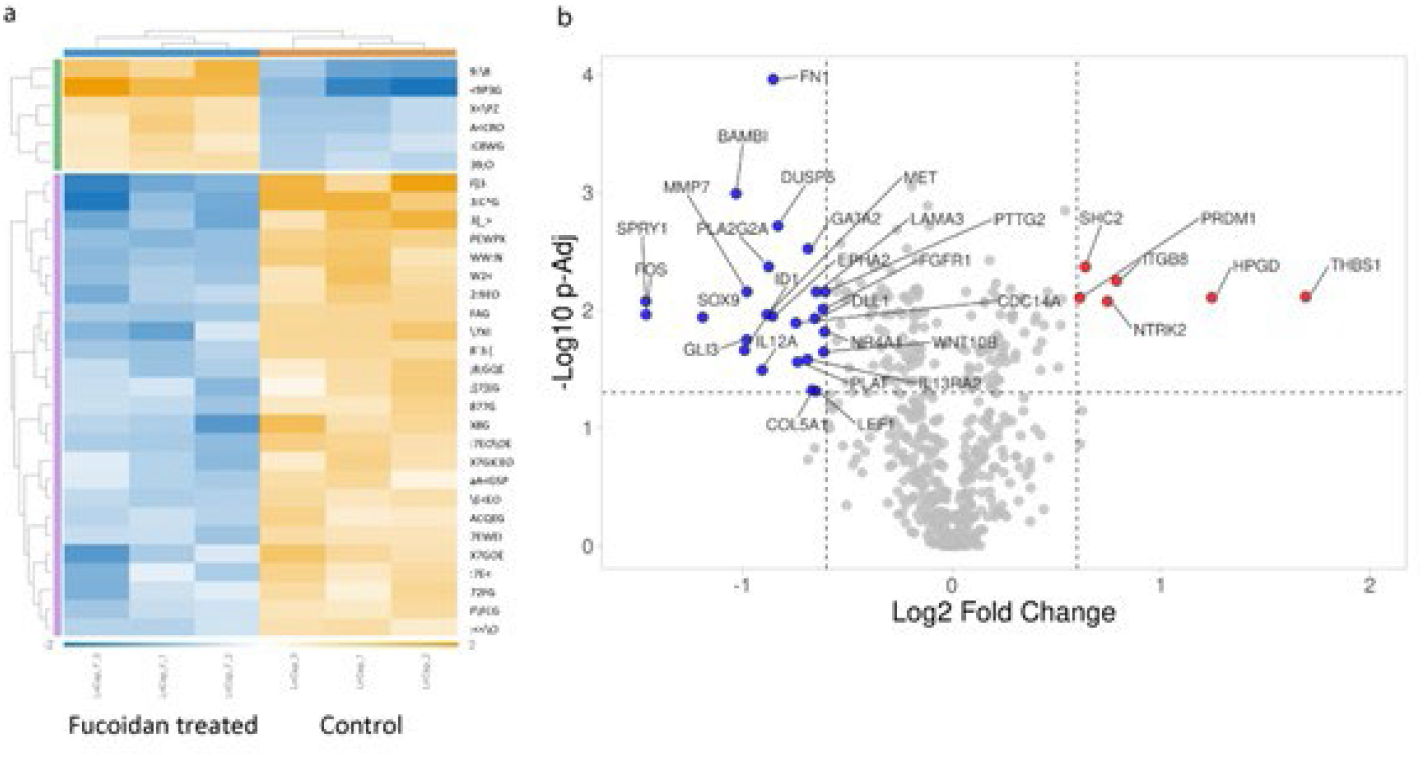
(a) A heatmap of differentially expressed genes between treated and untreated LNCaP cells. Each row is a gene and each column a cell line. Yellow is high expression and blue is low expression. (b)Volcano plot showing differentially expressed genes between LNCaP cells treated with fucoidan (LNCaP F) v untreated cells (LNCaP).

Pathway scores between the treated and untreated LNCaP spheroids indicates that treated cells were enriched in the MAPK, TGF-β, cell cycle/apoptosis, the JAK-STAT and Wnt pathways while the untreated cells were enriched in the PI3K-AKT, transcriptional misregulation, DNA Damage/Repair and Hedghog pathways (Figure 5).

**Figure 5.**
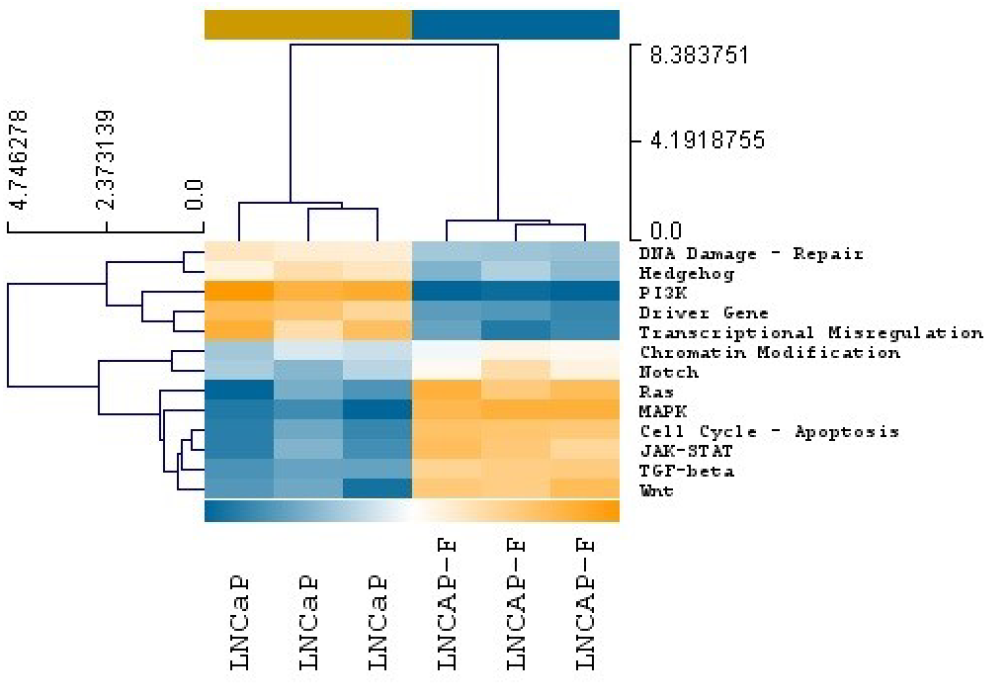
Gene set enrichment scores for LNCaP cells treated with fucoidan and untreated LNCaP cells. Each column is a sample and each row a pathway. Red is increased pathway expression and blue is decreased pathway expression.

### MiRNA profiling

Of the 798 miRNAs, only hsa-miR-1246 was significantly different at an adjusted *p*-value between the fucoidan treated LNCaP and non-treated cells, with 28 miRNAs exhibiting a statistical trend (*p*<0.1) for a difference between the conditions (Figure 6c). hsa-miR-1246 exhibited a statistical trend for a difference between the fucoidan treated PC-3 cells and the non-treated PC-3 spheroids (Figure 6b and Supplementary Table 3 and Supplementary Table 4).

**Figure 6.**
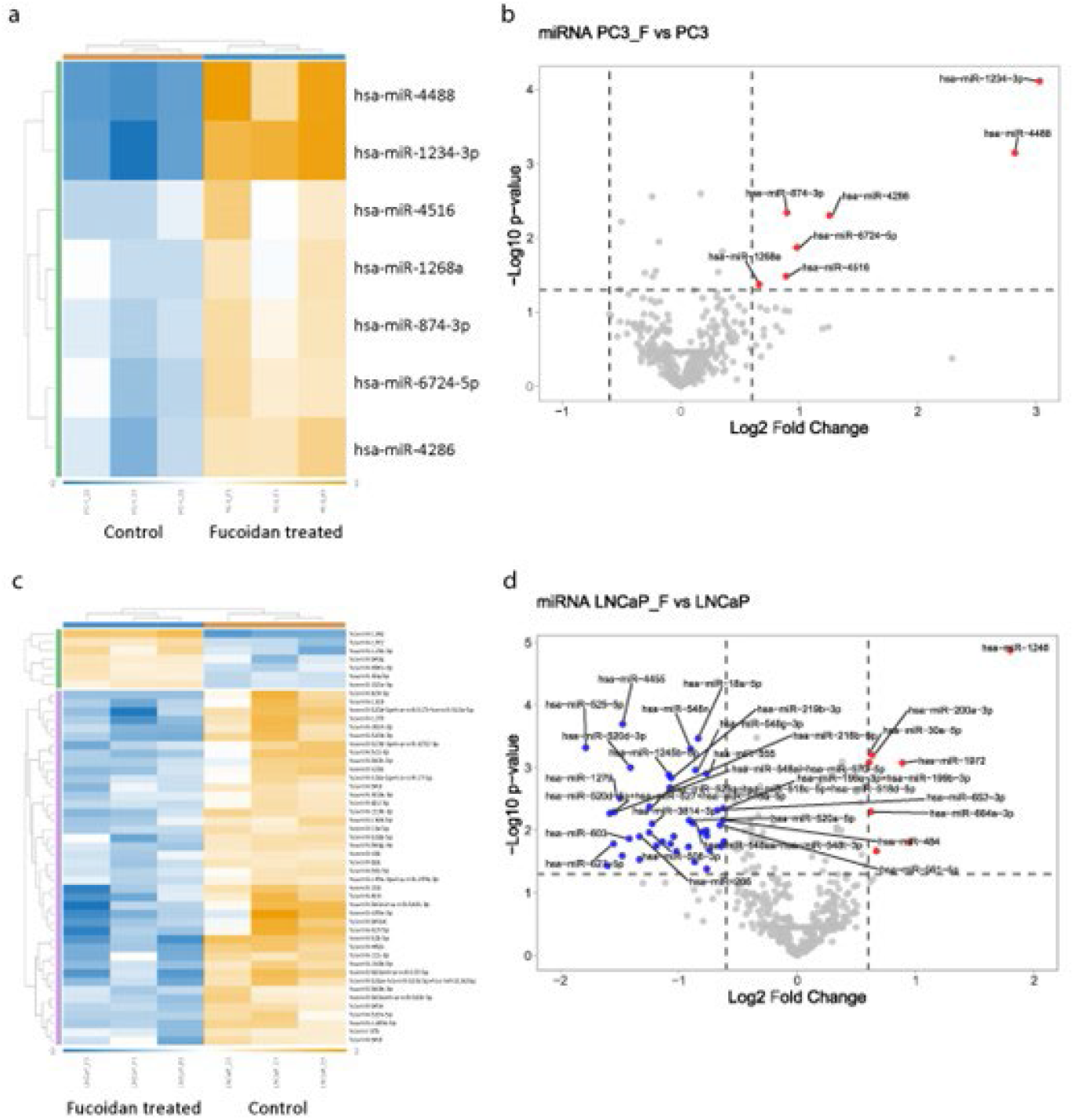
(a, c) Heatmaps of differentially expressed miRNAs between treated and untreated PC-3 and LNCaP cells. Each row is a gene and each column a cell line. Yellow is high expression and blue is low expression. (b,d) Volcano plots showing differentially expressed miRNAs between PC-3 and LNCaP cells treated with fucoidan (PC-3 F /LNCaP_F/) v untreated cells (PC-3/ LNCaP).

## Discussion

In this study, a proprietary fucoidan blend from *Undaria pinnatifida* and *Fucus vesiculosus* was evaluated for anti-cancer effects in 3D spheroid cultures of prostate cancer cells. Two well established prostate cancer in vitro cell models, a hormone insensitive cell line, PC-3 and a hormone sensitive cell line, LNCaP were assessed for changes in cell viability, spheroid structure, mRNA and miRNA expression. Fucoidan concentrations above 6.25µg/ml were observed to have a negative impact on cell viability and spheroid integrity in the PC-3 cell line, correlating with previous 2D culture studies, which had shown impacts on cell viability from 10 µg/ml [9].The disruption of the compact spheroid phenotype observed in the PC-3 cell line following exposure to the fucoidan blend may be related to cytotoxicity and cell apoptosis given data from the LDH assay aligned closely with the loss of spheroid structure and there was an enrichment of genes in the apoptosis pathway. However, direct interaction by fucoidan with pathways related to cell-cell adhesion cannot be ruled out. A depletion of genes in several cancer development and proliferation pathways was also observed in the PC-3 cells exposed to fucoidan.

Interestingly, a cytotoxic response was only observed at the top concentration (>100 µg/ml) for the LNCaP cell line likely confined to the outer cell layers, as commonly seen in dense cancer spheroid models. No impact on LNCaP spheroid structural integrity was observed up to the top dose of the assay (100 µg/ml). There was an enrichment of genes in apoptosis pathways observed in treated LNCaP spheroids, with pathways related to cancer development unchanged with treatment. Interestingly, fucoidan treated LNCaP cells had an increased expression of the microRNA hsa-miR-1246, a recognised tumour suppressor in prostate cancer. The changes in spheroid morphology and compound resistance described here illustrate that 3D cell culture models are an important tool for assessing the potential anti-cancer activity of compounds with novel modes of action. Fucoidan may be a promising natural product to target pathways that play a role in prostate cancer development and drug resistance.

Spheroid 3D cultures have several advantages over monolayer cultures in that they allow for assessment of intracellular interactions between cells and particularly anti-cancer agents and tumour cells. Clonal growth of tumour cells and spheroid morphology can also be included in biomarker identification and to determine the effect of treatment (27). A notable difference was observed between the two cell lines in the dosage of fucoidan required to alter spheroid formation, migration and viability. Cancer spheroid formation and cellular compaction relies on cell-cell adhesion and contraction (28). It may be that the loss of PC-3 spheroid structure was mediated by direct interaction of fucoidan in cell-cell adhesion. Previous reports have noted that fucoidan inhibits and blocks adhesion of other cancer cell lines to tissue substrata and other cells (29, 30). PC-3 spheroid disintegration due to the loss of cell-cell adhesion may also explain the cytotoxic effect of fucoidan observed through increased LDH concentrations. In contrast, the resistance of the LNCaP spheroid to disintegration may have hampered penetration of fucoidan into the spheroid regions and maintained cell survival. The increased LDH from LNCaP cells only at higher concentrations of fucoidan was likely released by the spheroid outer cell layers, as has previously been described (27, 31). The direct cytotoxic effects of Fucoidan on PC-3 cells make it a promising strategy to investigate clinically particularly in the context that serum uptake is very low and there is no information to the authors’ knowledge of how long serum levels persist following ingestion of fucoidan in supplemental form. Given that PC-3 cells are considered as a model of prostatic small cell neuroendocrine carcinoma (SCNC), a highly aggressive tumour, the sensitization of PC-3 spheroids by fucoidan indicates that it may also have promise as an adjuvant strategy to reduce SCNC drug resistance.

Gene expression profiling revealed several potential mechanisms by which the fucoidan blend may have prostate anti-cancer effects. Compared to the non-treated cells, fucoidan exposure inhibited the PI3K pathway and increased cell cycle apoptosis pathways in both hormone dependent and hormone independent cells. Several studies strongly implicate the over-activation of the PI3K-AKT-mTOR pathway in prostate cancer progression (32-34). Fucoidan blend incubation also induced notable differences between the two cells lines in gene expression in other oncogenic pathways. In the hormone independent PC-3 cell line, changes in gene expression indicate an inhibition of inflammatory pathways, in particular the PI3K, Ras, Wnt and JAK-STAT pathways compared to the hormone dependent LNCaP cell line, in which there was increased gene expression in both anti-cancer and pro-cancerous pathways. It may be that the use of anti-inflammatory agents that target the MAPK and JAK-STAT pathways in conjunction with fucoidan should be considered for hormone dependent prostate cancer investigations. The biology underpinning tumour initiation and progression involves a complex interplay between several physiological systems and this study demonstrates that fucoidan blend is able to directly modulate key anti-cancer pathways.

Our observation that the fucoidan blend did not have a greater impact on miRNA expression in either hormone dependent or independent cell lines was surprising. miRNAs function through mRNA degradation or post-translational inhibition and circulating miRNAs have shown promise for prostate cancer diagnosis either alone (35) or in combination with prostate serum antigen (36), and with treatment outcomes (37). Furthermore, downregulation of hs-miR-141-3p has been associated with prostate cancer tumour progression (38). Interestingly, treatment of LNCaP cells with the fucoidan blend was associated with an increase in hsa-miR-1246, a miRNA that has been shown to increase apoptosis and decrease cell proliferation in previous prostate cancer research (39). *In-vitro* and *in-vivo* hsa-miR-1246 expression has been found to be low in prostate cancer and is considered a potent tumour suppressor via indirect regulation of the PI3K/AKT signaling pathway (40). No other established miRNAs, such as hs-miR-375, hsa-miR-92a-3p or let-7 had altered expression in either cell line to the fucoidan blend.

Dietary inventions for the prevention and attention of prostate cancer have been discussed by others (41, 42). Various dietary additions ranging including potato extracts (43) and citrus extracts (44) have been examined *in vitro, in vivo* and in clinical trials (45) for their potential as modulators of disease progression. Whilst there are no current comparative data, the low concentrations needed to exert *in vitro* effects on spheroid formation and key tumourigenic pathways in PC-3 cells and on hsa-miR-1246 expression in the hormone dependent LNCaP cells, suggest that fucoidan may also be a promising dietary intervention for prostate cancer. In a previous study examining the effects of fucoidan supplementation and circulating miRNA expression, ingestion of a single dose of *Undaria pinnatifida* fucoidan by healthy normal subjects was associated with changes across 29 pathways, including those associated with immunity, cancer cells, inflammation, and neurological function (46). This clearly indicates the ability of fucoidan to alter the miRNA environment in normal subjects and indicates that changes in miRNA may also be possible in prostate cancer patients. Future clinical research examining whether oral fucoidan can alter mRNA or miRNA in prostate cancer is now essential, particularly given the continued rate of 5-year survival with this disease remains at ∼30% (47).

This study further supports the use of 3D spheroid cultures to gain a more nuanced understanding of the way in which different cell types respond to treatment. Based on the data shown here, we hypothesize that ingestion of fucoidan exerts cytotoxic effects on hormone resistant prostate cancer and may alter blood mRNA and miRNA expression in both hormone sensitive and hormone resistant prostate cancer. Future research should consider assessment with more complex in vitro modelling including patient-derived tumor spheroids that cover more tumour heterogeneity and in vivo models that may assess potential immune system modulation. Quantifying corresponding cell-cycle or DNA damage-related protein levels would also assist in establishing potential fucoidan mode of action. Ultimately clinical assessment in patients with diagnosed prostate cancer before and after dietary supplementation with fucoidan could be undertaken.

## Conclusions

The Fucoidan blend examined in this study specifically disrupted spheroid integrity and exerted cytotoxic effects on the hormone resistant PC-3 cell line. Identifying potential anti-cancer properties of marine algae such as the fucoidan blend examined here has traditionally been performed in standard monoculture conditions. The use of 3D spheroid culture models has extended the value of cell line research techniques for compound assessment in prostate cancer research (48). Utilising gene expression profiling of spheroid cultures, we found that fucoidan treatment of hormone sensitive LNCaP cells and hormone resistant PC-3 prostate cancer cells altered several pathways involved in cancer progression. Prominent amongst these is inhibition of the PI3K-AKT pathway. The PI3K-AKT pathway is overactivated in prostate cancer and is associated with prostate cancer progression and PI3K inhibitors have demonstrated anti-prostate cancer results. Fucoidan also increased the expression of the prostate tumour suppressor hsa-MiR-1246 in hormone sensitive cells. These findings provide important pre-clinical evidence for further studies into therapeutic applications of fucoidan in prostate cancer.

## Abbreviations

Pca: prostate cancer
iRNA: microRNA
mRNA: messenger RNA
LNCaP: Lymph node carcinoma of the prostate
PC-3: prostate cancer cell line
FBS: fetal bovine serum
LDH: lactate dehydrogenase
FOV: field of view
SD: standard deviation
CI: confidence interval
µ: micro
mg: milligrams

## Disclosure

### Data availability Statement

Data is available through the corresponding author.

### Ethics Statement

-Approval of the research protocol by an Institutional Reviewer Board: N/A

-Informed Consent: N/A

-Registry and the Registration No. of the study/trial: N/A

-Animal Studies: N/A

### List of Supporting information

**Supplementary Figure 1. LDH release from PC-3 and LNCaP cells cultured in the presence of the fucoidan blend. Data normalized to top concentration dose effect**.

**Supplementary Table (ST) 1. Differentially expressed genes between PC-3 cells treated with fucoidan and untreated PC-3 cells**

**ST2. Differentially expressed genes between LNCaP cells treated with fucoidan and untreated LNCaP cells**.

**ST 3. Differentially expressed miRNA between fucoidan-treated LNCaP cells and untreated cells**.

**ST4. Differentially expressed miRNA between fucoidan-treated PC3 cells and untreated PC-3 cells**

