## Supplementary Figures for "Fucoidan exerts anti-cancer activity in a 3D prostate spheroid cell culture model"

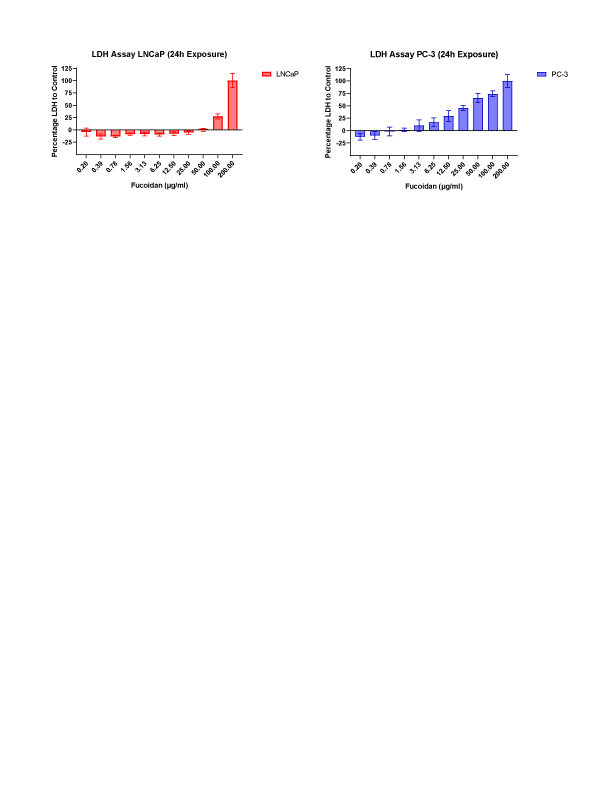

SF1. LDH release from PC-3 and LNCaP cells cultured in the presence of the fucoidan blend. Data normalized to top concentration dose effect.

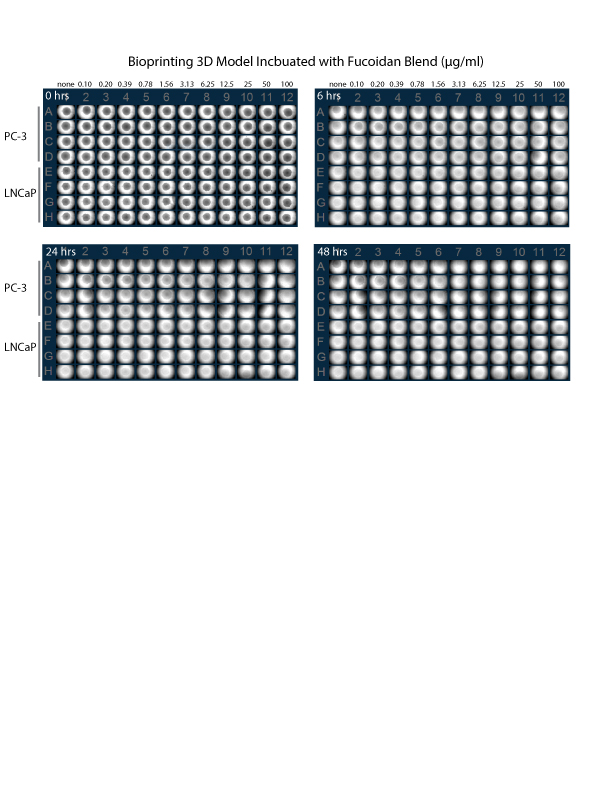

SF2. Representative 3D culture overview in plate format for PC-3 and LNCaP cell lines at 0, 6, 24, 48-hour timepoints.

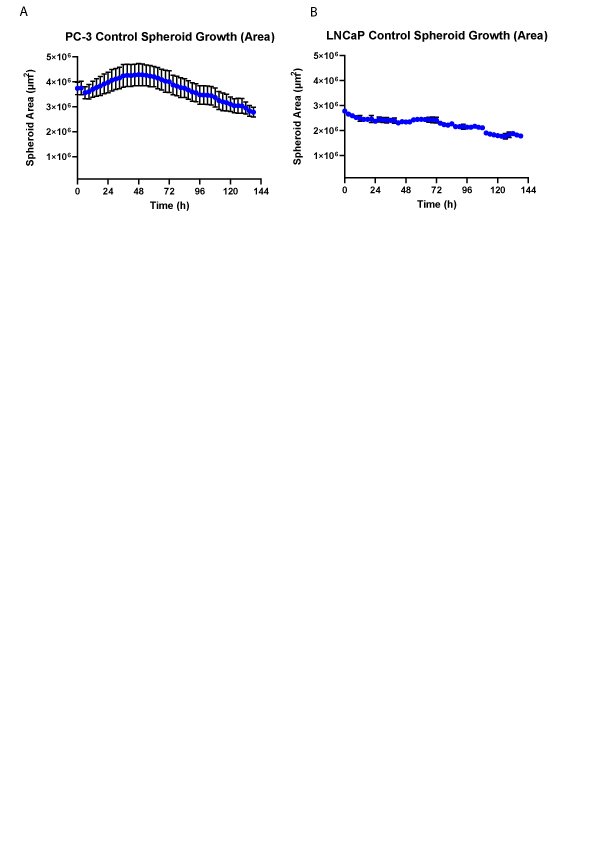

**SF3.** (A) PC-3 and (B) LNCaP spheroid area from vehicle control wells over time (30min datapoints captured on an Incucycte live cell analysis System). Data points represent quadruplicate spheroids +/-SD.

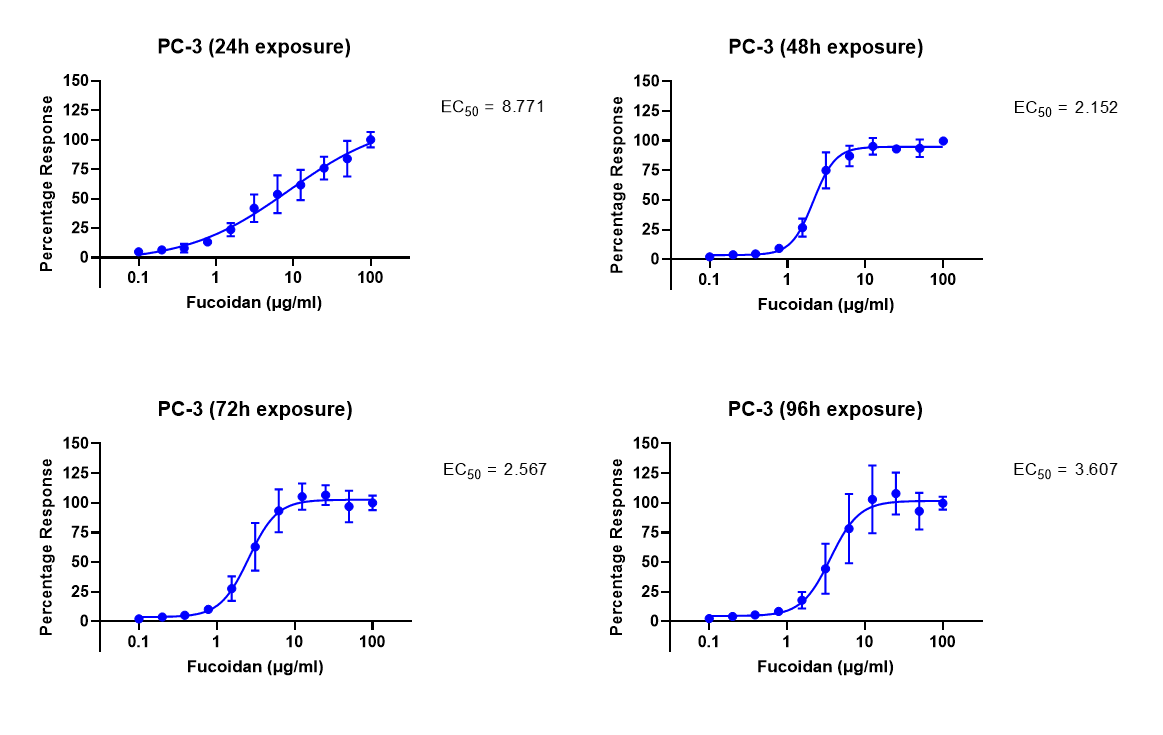

SF4 Additional EC_50_ time point data from PC-3 spheroid area measurement (24h – 96h). Average of quadruplicate data points with SD error bars.

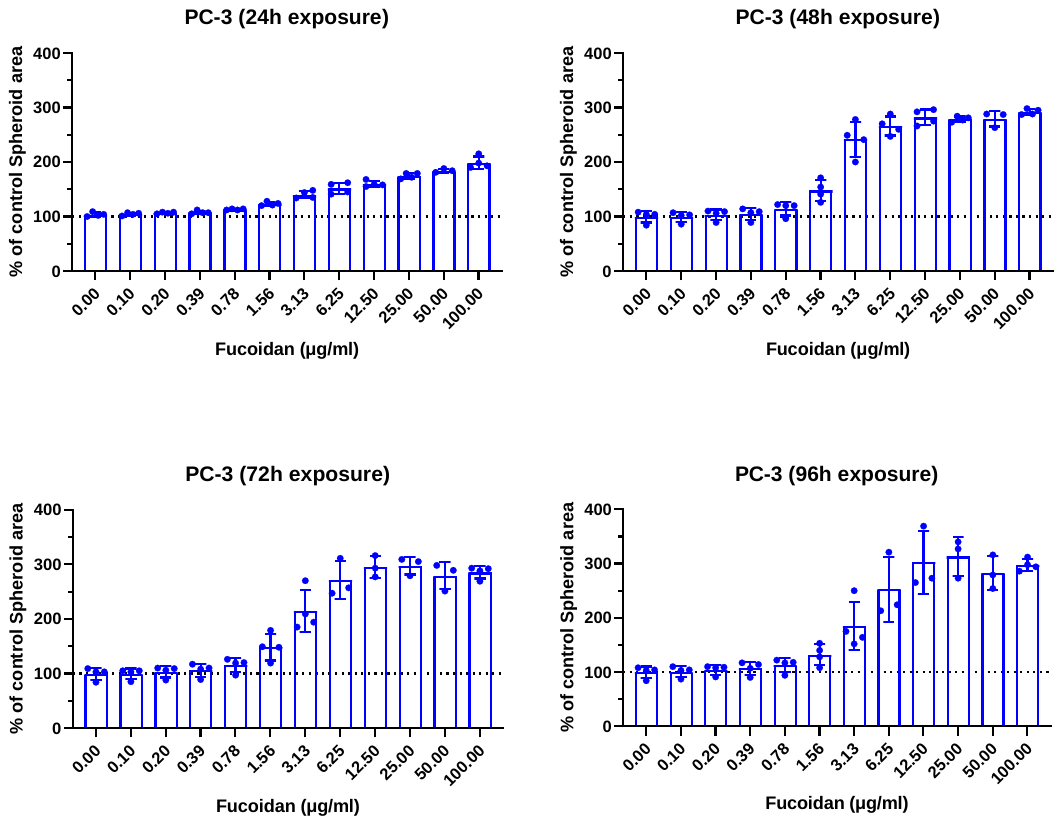

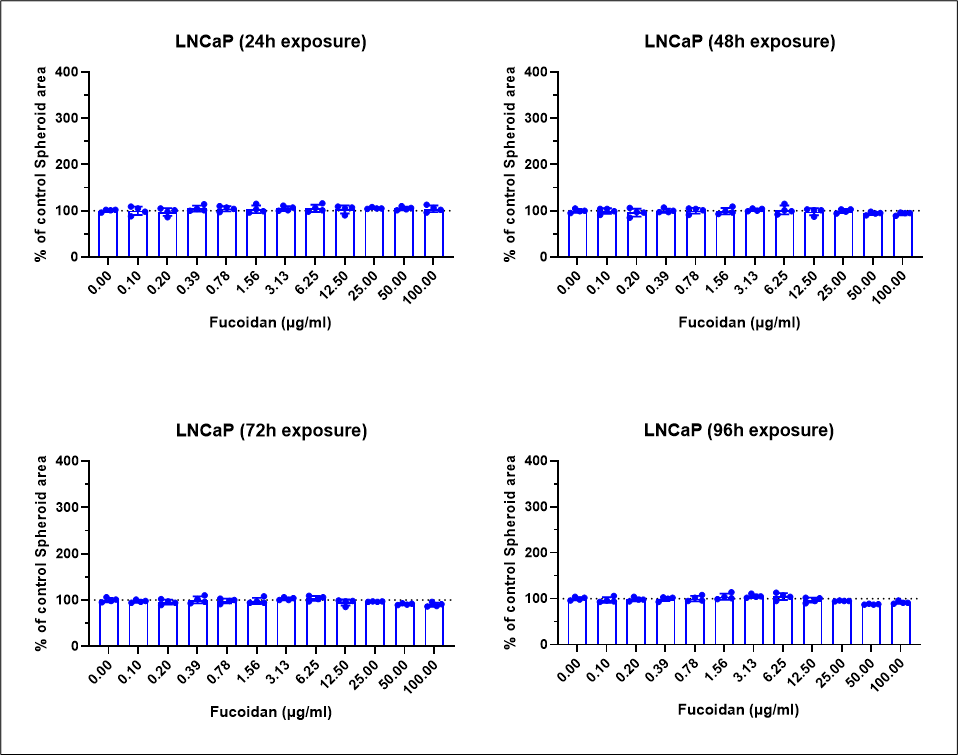

SF5. Additional timepoint spheroid area data for both PC-3 and LNCaP cell lines (24h – 96h). Average of quadruplicate data points with SD error bars.
